# Cleaner wrasse pass the mark test. What are the implications for consciousness and self-awareness testing in animals?

**DOI:** 10.1101/397067

**Authors:** Masanori Kohda, Takashi Hotta, Tomohiro Takeyama, Satoshi Awata, Hirokazu Tanaka, Jun-ya Asai, L. Alex Jordan

## Abstract

The ability to perceive and recognise a reflected mirror image as self (mirror self-recognition, MSR) is considered a hallmark of cognition across species. Although MSR has been reported in mammals and birds, it is not known to occur in any other major taxon. A factor potentially limiting the ability to test for MSR is that the established assay for MSR, the mark test, shows an interpretation bias towards animals with the dexterity (or limbs) required to touch a mark. Here, we show that the cleaner wrasse fish, *Labroides dimidiatus*, passes through all phases of the mark test: (*i*) social reactions towards the reflection, (*ii*) repeated idiosyncratic behaviours towards the mirror (contingency testing), and (*iii*) frequent observation of their reflection. When subsequently provided with a coloured tag, individuals attempt to remove the mark in the presence of a mirror but show no response towards transparent marks, or to coloured marks in the absence of a mirror. This remarkable finding presents a challenge to our interpretation of the mark test – do we accept that these behavioural responses in the mark test, which are taken as evidence of self-recognition in other species, mean that fish are self-aware? Or do we conclude that these behavioural patterns have a basis in a cognitive process other than self-recognition? If the former, what does this mean for our understanding of animal intelligence? If the latter, what does this mean for our application and interpretation of the mark test as a metric for animal cognitive abilities?

## Introduction

The mark test, in which a coloured mark is placed on a test subject in a location that can only be viewed in a mirror reflection, is held as the benchmark behavioural assay for assessing whether an individual has the capacity for self-recognition [1,2]. In human infants, approximately 65% of individuals pass the mark test by 18 months of age by touching the mark with their hands while viewing their reflection [3], although some individuals pass earlier and some never pass. Accumulating reports claim that many other animal species also pass the mark test, including chimpanzees [1], elephants [4], dolphins [5,6] and corvids [7], while many other species are apparently unable to pass the test [8; but see 9-11]. Nevertheless, the interpretation of these results is subject to wide debate, and the certainty with which behavioural responses during the mirror test can be taken as evidence of self-awareness in these animals is questioned (8,12,13). This problem is exacerbated when the taxonomic distance increases between the test species and taxa for which the test was initially designed. Can for example the behavioural results recorded for primates during the mirror test be meaningfully compared with those in birds? If yes, does this mean a bird that passes the mirror test is self-aware? More generally, if we are interested in understanding and comparing cognition and problem-solving across taxa, can we assume that equivalent behaviours represent equivalent underlying cognitive processes? With particular reference to the mark test, here we explore what forms of behaviour in fish could be taken as evidence of self-awareness, as has been done for primates and other taxa.

Given that the mark test as designed for humans and primates relies on hand gestures toward the marked region and changes in facial expression, we also ask whether it is even possible to interpret the behaviour of ‘lower’ taxonomic groups during the mark test in the same way as for ‘higher’ taxa. If not, we must ask how useful the mark test is as a test for self-awareness in animals. To explore these questions, we here test whether a fish, the cleaner wrasse *Labroides dimidiatus*, displays behavioural responses that constitute passing the mark test. We then ask what this may mean for our understanding of self-awareness in animals and our interpretation of the test itself.

To date, no taxon outside of birds and mammals has passed the mark test. This is despite many species in other vertebrate classes, such as fish, showing sophisticated cognitive capacities in other tasks [14-17], including transitive inference [18,19], episodic-like memory [20], playing [21], tool use [22,23], prediction of the behaviour of others by using one’s own experience during coordinated hunting [24, 25], cooperating to warn about predators [26,27] and cooperative foraging [28]. These studies reveal that the perceptual and cognitive abilities of fish often match or exceed those of other vertebrates [15,17], and suggest the possibility that the cognitive skills of fish could more closely approach those found in humans and apes [14,16,17,24,28].

Nevertheless, it can be challenging to employ standardised cognitive tests across species when performance in the test depends on specific behavioural responses that are not present in all taxa. This may be considered the case for the mark test, which has been designed to suit the behavioural repertoire of humans and primates [1,2]. Animals that cannot directly touch the marks used in MSR tests, such as fish as well as dolphin, are therefore regarded as poor test candidates [2,5,29], making direct comparison of their cognitive capacities with those of other vertebrates challenging [30-33]. Although no mark tests were performed, behavioural observation of manta rays (Chondrichthyes) on exposure to a mirror suggests that these fish show self-directed behaviour toward their reflection [34], though these results are contested [35,36]. This controversy highlights the need to question what type of behavioural response would be taken as evidence of interacting with the mark in an animal with the morphology of a ray, and whether these behaviours may have an alternative function, e.g. in social communication. To make a comparison across taxa, one must carefully consider the inherent biology of the focal species, preferably choosing a species with perceptual abilities and a behavioural repertoire that i) allow it to respond to coloured marks placed on the body (this is not a given when the sensory systems of animals differ so greatly) and ii) do so in a manner that can be interpreted by a human observer.

The cleaner wrasse, *Labroides dimidiatus*, is such a species, forming mutualistic relationships with larger client fish by feeding on visually detected ectoparasites living on the skin of the clients [37]. Therefore, the cleaner wrasse has sensory and cognitive systems that are well-equipped for visually detecting spots of unusual colour on the skin surface, as well as the behavioural repertoire required to respond to marks. This species is highly social, interacting with the same individuals repeatedly over long periods of time, and has sophisticated cognitive abilities, including tactical deception [38-40] and reconciliation [41], and it can also predict the actions of other individuals [41,42]; these are traits requiring mental abilities that may be correlated with the ability for self-recognition, as seen in other species [16,29,43-45].

During the mirror test, animals must visually locate a mark in a mirror image. The interpretation of the test is that animals regard the mark on the body as unusual and thus examine it. It is reasonable to predict the wrasse will perceive the coloured marks as visually similar to ectoparasites, and may thus evoke an attentional response that may culminate in a removal attempt [46-47]. However, lacking hands and arms, any attempt to remove or interact with the mark would necessarily take a different form to that observed for primates or elephants. Fortunately for the question at hand, many fish taxa, including cleaner wrasse, display a characteristic behaviour that functions to remove irritants and/or ectoparasites from the skin surface [48,49], termed glancing or scraping. This behaviour may therefore be considered as self-directed, just as for some mammal species that also lack hands in which scraping is taken as an indicator of self-directed behaviour during the mirror test [29,50]. We therefore consider the cleaner wrasse to possess the necessary sensory biology and behavioural repertoire to adequately employ the mirror test, and here use a modified experimental paradigm established for studies of humans and apes to test for mirror self-recognition in a fish. Importantly, this species allows us to ask whether the criteria that are accepted as evidence for mirror self-recognition in mammals and birds can be applied to other taxa, and if they fulfil these criteria, what it means for our interpretation of the test itself.

In applying the mirror test, transitions among three behavioural phases after initial exposure to a mirror are typically [1,4,5,6]; these transitions among behavioural phases are interpreted as additional evidence of self-recognition, although in themselves do not constitute passing the mirror test [1,4]. We first tested whether the cleaner wrasse passed through all three behavioural phases upon exposure to a mirror placed in an experimental tank (Fig. 1A), and if so, we describe the phases in cleaner fish. The first phase (*i*) is a social reaction towards the mirror, apparently as a consequence of the reflection being perceived as an unknown conspecific. In phase (*ii*), animals begin to repetitively perform idiosyncratic behaviours that are rarely observed in the absence of the mirror. These behaviours are interpreted as contingency testing between their own actions and the behaviour of the reflection [e.g. 1,4]. In phase (*iii*), the animal begins to gaze and examine their reflection as if it is a representation of the self, and uses the mirror to explore their own body in the absence of aggression and mirror-testing behaviour [1,4,5]. If they passed these phases, we applied the mark test.

**Fig. 1.**
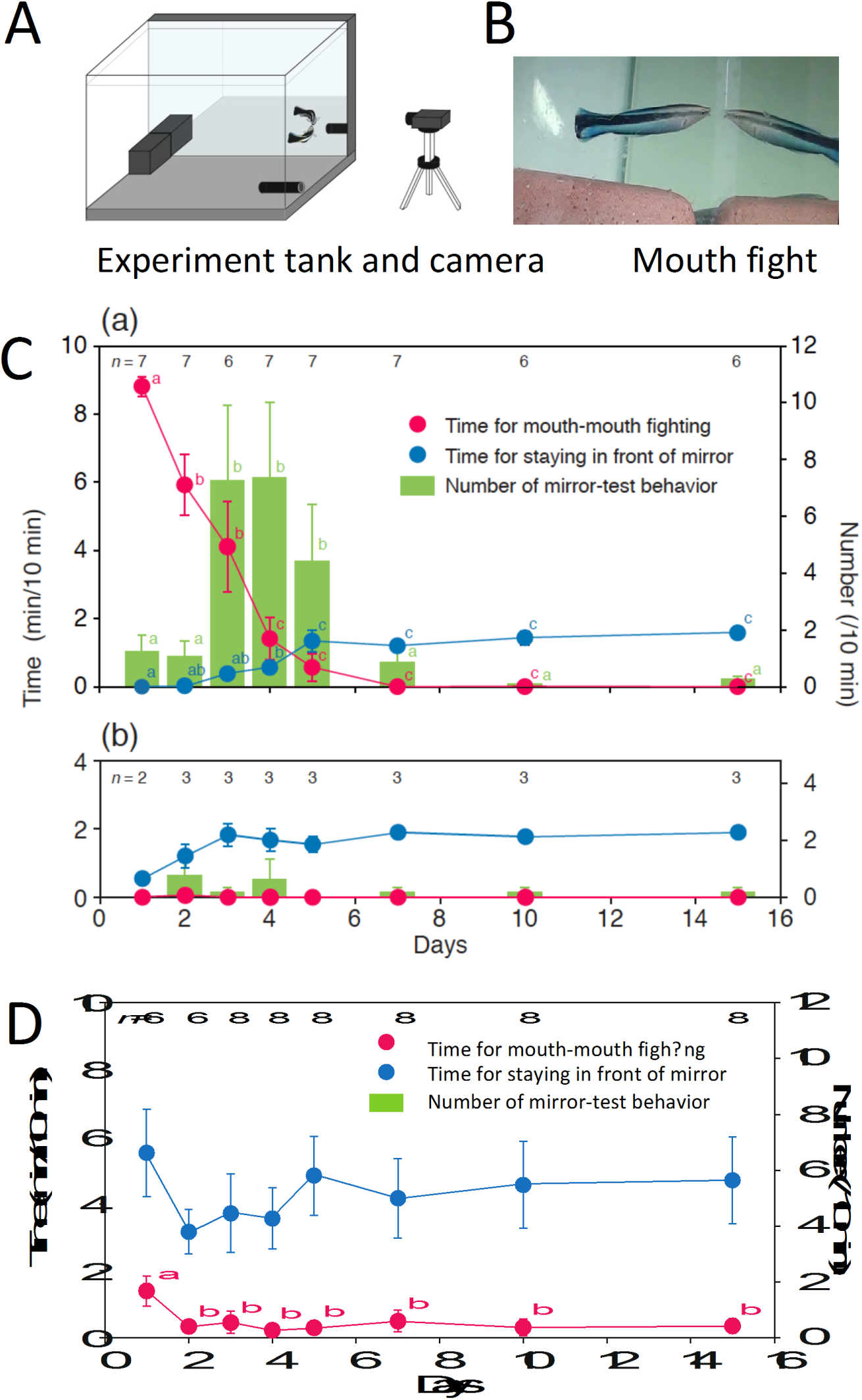
Responses of cleaner wrasse to the mirror and to real fish during the 2-week period after the mirror was introduced into the experimental tank. (A) Design of the experimental tank showing the mirror location. (B) Photograph of mouth-to-mouth fighting against a mirror reflection. (C) Change in social responses towards the mirror. Mean ± SE for time spent mouth fighting (red), time spent within 5 cm of the mirror without being aggressive (blue), and frequency of mirror-testing behaviours/10 min (green). Superscript labels a, b and c denote statistical differences. C1: on the seven fish appearing in Table 2; C2: on the three fish #4, #5 and #6. Statistical results for daily changes in time spent mouth fighting, LMM, c_7_^2^ = 91.87, *P* < 0.0001; time spent in front of the mirror, LMM, c_7_^2^ = 64.63, *P* < 0.0001, and changes in the number of mirror-testing behaviours, GLMM, c_7_^2^ = 137.08, *P* < 0.0001. (D) Change in social responses to live conspecific fish over 2 weeks: Statistical results for daily changes in time spent mouth fighting, LMM, df = 7, chi-square = 27.36, *P* = 0.0003, and time spent in front of the mirror, LMM, df = 7, chi-square = 9.09, *P* = 0.25; no idiosyncratic behaviours were observed in any fish in this condition. Symbols are as in (C).

## Results and discussion

### Progression of behaviours in response to the mirror

Prior to starting the experiments, the focal fish swam around the tank and showed no unusual reactions to the covered mirror. Immediately after initial exposure to the mirror, 7 of 10 fish responded aggressively to their reflection, attacking it and exhibiting mouth-to-mouth fighting [45,46] (Fig. 1A,B, Supplementary Movie S1) suggesting that the focal fish viewed the reflection as a conspecific rival. The frequency of mouth fighting was highest on day 1,and decreased rapidly thereafter, with zero occurrences by day 7 (Fig. 1Ca; cf. with the similar decrease in aggression seen in chimpanzees, and shown in Fig. 2 of [1]) and hardly any aggression over the following month.

**Table 2.** Number of atypical (mirror-testing) behaviours shown by seven fish during the 20-min observation period in the first 5 days after presenting the mirror. See Table 1 for description of the behaviours. (*a*): Dashing along the mirror, (*b*): Dashing with head in contact with the mirror, (*c*): Dashing and stopping, (*d*): Upside-down approach, (*e*) Quick dance The most frequent mirror-testing behaviour of each fish is underlined.

| 880 Fish |  | Categories of mirror-testing behaviour |  |  |  |  |  |
| --- | --- | --- | --- | --- | --- | --- | --- |
|  |  | (a) | (b) | (c) | (d) | (e) | Total |
| 882 | #1 | 2 | 0 | 4 | <u>39</u> | 0 | 45 |
|  | #2 | 0 | 0 | <u>30</u> | 0 | 0 | 30 |
| 884 | #3 | <u>54</u> | 0 | 0 | 0 | 0 | 54 |
|  | #7 | 15 | 0 | 0 | 0 | <u>35</u> | 50 |
| 886 | #13 | <u>33</u> | 0 | 0 | 0 | 0 | 33 |
|  | #20 | <u>31</u> | 17 | 2 | 0 | 2 | 52 |
| 888 | #21 | 2 | 0 | 2 | 0 | <u>6</u> | 10 |
|  | Total | 137 | 17 | 38 | 39 | 43 | 274 |

**Fig. 2.**
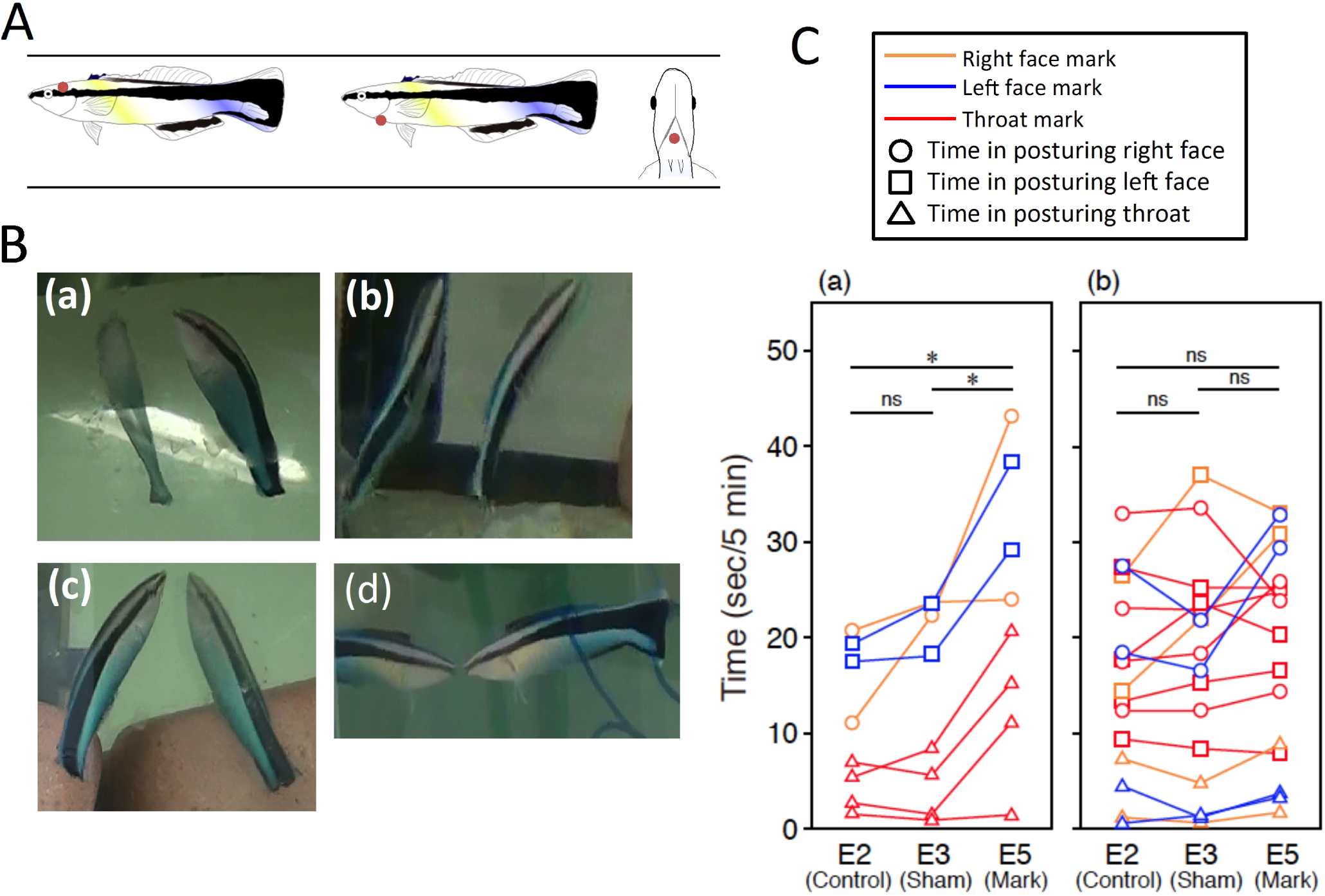
Mark locations and time spent in a posture allowing viewing of the marked site. (A) Marked location on the left side of the head (left) and under the throat (right); (B) Postures performed in front of the mirror: (a) displaying right side to mirror, (b) displaying left side to mirror, (c) vertical posture displaying throat, and (d) frontal-vertical posture. (C) Duration of each posture performed in the mirror for each individual: fish marked on the right side of the head (yellow, *n* = 2), left side of the head (blue, *n* = 2), and throat (red, *n* = 4) during periods E2 (control: no mark), E3 (sham-mark) and E5 (coloured mark with mirror). (C-a): Time spent viewing the marked location (i.e. the correct side): Repeated-measures ANOVA, main effect of sequences: *F* = 12.09, *P* = 0.016, marked position: *F* = 19.06, *P* = 0.005, sequence × marked position: *F* = 0.70, *P* = 0.54. * < 0.05, ns = not significant (*n* = 8) (C-b): Time spent viewing unmarked positions. Repeated-measures ANOVA, sequence: *F* = 2.54, *P* = 0.12, marked position: *F* = 13.15, *P* = 0.0008, sequence × marked position: *F* = 0.99, *P* = 0.42. The posture allowing head viewing was performed more frequently than the vertical posture, because vertical swimming is rare in this species.

As mouth fighting towards the mirror reflection decreased, the incidence of unusual and atypical behaviours (e.g. ‘upside-down approach’ and ‘dashing along mirror’; Table 1, Movies S2, S3) significantly increased and was highest on days 3–5 (Fig. 1Ca). On days 3 and 4, the estimated average frequency of these atypical behaviours among the seven individuals was 36 times per hour during the daytime. Each of these atypical behaviour types was of short duration (≤1 s), often consisting of rapid actions that occurred suddenly within 5 cm of the mirror. At the end of each movement, the fish remained near the mirror, and appeared as if they were viewing their reflection (Movies S2, S3). These atypical behaviours could be loosely grouped into five types: not against the mirror reflection, dashing along the mirror without and with attaching the head on the surface (atypical behaviours, *a* and *b, respectively*); dashing towards the reflection but stopping before touching it (*c*); and idiosyncratic postures and actions of short duration performed in front of the mirror: upside-down approach (*d*), and quick dance (*e*) (Table 1). While it is possible to interpret these behaviours as a different form of aggression or social communication, they have not been recorded in previous studies of social behaviour in this species [46] and were not part of a courtship display, as all of the subject fish were females.

These atypical behaviours were highly repeatable within an individual, with each fish performing one or two types of behaviour more than 400 times a day on average during days 3 and 4 (Table 2; Fisher’s exact probability test for count data with simulated *P*-value based on 2,000 replicates of *P* = 0.0005). Crucially, these behaviours occurred only upon exposure to the mirror, and were not observed in the absence of the mirror (i.e. before mirror presentation). Almost all of the behaviours ceased by day 10 (Fig. 1Ca), and thereafter were hardly observed at all over the following month.

These behaviours are different from the previously documented contingency-testing behaviours of great apes, elephants and magpies [1,4,7], but given the taxonomic distance between them, this could hardly be otherwise. While primates and elephants may perform more anthropomorphic behaviours such as changing facial expression, or moving the hands, legs or trunk in front of the mirror, wrasse and other fishes cannot perform behaviour that is so easily interpreted by a human observer. Nevertheless, behaviours such as rapid swimming and other spontaneous actions could represent alternative indices of contingency that are within the behavioural repertoire of the study species (Table 1).

In summary, the atypical movements observed in cleaner wrasse were characterised by almost every aspect of contingency-testing behaviour documented previously [1,4,5,7]: 1) atypical and idiosyncratic, 2) occurring repeatedly, 3) occurring in front of a mirror, 4) not occurring in the absence of a mirror, 5) occurring after a phase of initial social behaviour, 6) occurring over a short period of time and 7) distinct from aggressive behaviour. Fulfilment of these conditions supports the contingency-testing hypothesis. Although we reserve judgement as to whether these behaviours should be interpreted as evidence that the fish examine and perceive the reflection as a representation of self, we nevertheless conclude that these behaviours are consistent with phase (*ii*) of MSR as presented for other taxa.

In phase (*iii*), which is difficult to clearly distinguish from phase (*ii*), species that pass the mark test increase the amount of time spent in front of the mirror in non-aggressive postures while viewing the mirror image [1,4,5,7]. This interpretation is again rife with pitfalls, as it requires an assessment of the intentionality of unusual animal behaviours. An agnostic approach is to simply measure the amount of time animals spend in postures that reflect the body in the mirror [2]. This gives an upper measurement of the time in which animals could be observing their reflection while making no inferences about the intentionality of the act. We observed an increase in the amount of time spent in non-aggressive postures while close to the mirror (distance of < 5 cm), peaking on day 5 after mirror presentation and remaining consistently elevated (Fig. 1Ca; 107.0 sec ± 21.2 [SD]/10 min) versus days 1–4 and the several days prior to mirror presentation (37.0 sec ± 11.5, Wilcoxon sign-ranked test, *T* = 36, *P* = 0.008); this behaviour was consistent with phase (*iii*) of MSR. We did not observe specific viewing behaviour that is seen in chimpanzees and elephants, e.g. trying to look at body parts, such as inside the mouth or between the legs. It is inherently difficult to distinguish such looking behaviours from other behaviours primarily because gaze direction could not be determined in this species. Technological developments that allow eye tracking in free-swimming fish may alleviate this difficulty in future studies.

Similar to other studies, not all individuals we tested passed through each phase of the test. After the initial presentation of the mirror, three fish (#4, #5, #6) showed low levels of aggression and rarely performed atypical behaviours during period E1 (Fig. 1Cb). Instead, these three individuals spent relatively longer periods in front of the mirror, as is typically observed during phase (*iii*) in other focal fish (Fig. 1Cb). By applying the same criteria as applied for other instances of the test, we conclude these fish failed the test. However, an alternative explanation is that these fish had already passed through the initial phases; at the start of the experiment, the glass wall on the opposite side of the mirror in the tanks of these three fish was slightly reflective due to differences in lighting in the room, and the focal fish were observed to occasionally remain in front of the glass wall. These observations suggest that these three fish may have already passed through phases (*i*) and (*ii*) during the acclimation phase before the start of experiment. As discussed below, these three fish exhibited good responses to the mark test.

Species with MSR distinguish their own reflection from real animals viewed behind glass [e.g. 29]. When we exposed naïve cleaner wrasse to conspecifics behind glass, we observed fundamentally different responses towards their mirror image. Aggressive behaviour frequency towards real fish was generally low, yet did not diminish appreciably during the 2-week testing period (Fig. 1D). Time spent within 5 cm of the glass in the presence of conspecifics was also higher than that in the presence of the mirror. Importantly, no atypical or idiosyncratic behaviour (i.e. contingency-testing) was exhibited towards conspecifics.

These behaviours were only observed upon exposure to the mirror.

### Mark test

In the second part of the experiment, we used a modified standard mark test protocol to assess reactions to visible or sham (clear or un-pigmented) coloured marks. We used subcutaneously injected elastomer (see Material and methods; Northwest Marine Technology Inc. (NMTI), Anacortes, WA, USA). The coloured marks are applied with a small amount of coloured gel using a fine needle, a procedure, which has been repeatedly shown not to affect fish behaviour [51-54, NMTI], and is widely used in fish behavioural studies. Moreover, use of both visible coloured and invisible sham marks directly controlled for any effects of application, including irritation or tactile sensations around the marking site. Nevertheless, the procedure was necessarily modified from terrestrial tests due to the requirements of provisioning marks in the aquatic environment, and on animals covered in a protective mucus coating. Because of this, it involved potentially higher tactile stimulation than e.g. paint marks on elephant skin. We controlled for these direct tactile effects with a sham-marking experimental phase, observing no effect (see below). Nevertheless, we must consider a recent study that showed visual-somatosensory training induced self-directed behaviour in rhesus monkeys [10,11], which could not be achieved through visual stimuli alone. Our study differs in that we do not provide direct somatic stimulation during the mirror test, and much evidence suggests that no further tactile stimulation is maintained by elastomer injections after the procedure. However, given the nature of the mark application, we cannot rule out that a combination of visual and tactile cues produces the behavioural responses we describe.

Fish were marked at night while under anaesthesia, and they swam normally early the next morning under a no-mirror condition (Kohda, pers. obs.). After the initial settlement period “E1” (i.e. the initial 2 weeks of phases *i–iii*), we evaluated behaviour during periods “E2” (no mark), “E3” (injection with invisible sham mark), “E4” (injection with coloured mark with no mirror present) and “E5” (coloured mark with mirror present) during a further 2-week period. The sham and coloured marks were applied on the right side of the head of two fish, on the left side of the head of two other fish and under the throat in a further four fish (Fig. 2A); these areas were only visible in the mirror. Each mark was in the form of a small brown dot resembling an ectoparasite.

#### 1) Viewing coloured marks in the mirror

We first examined whether fish were assuming postures in front of the mirror that would reflect the marked site by categorising all body postures performed within 5 cm of the mirror into three types: postures exposing the right side of the head to the mirror, postures exposing the left-side of the head, and frontal-vertical postures exposing the head, throat, and underside to the mirror. These postures would respectively reflect the right face mark, the left face mark, and the throat mark (Fig. 2B). We predicted that if fish were attempting to observe the coloured marks on body parts reflected in the mirror, they would assume postures that facilitated this observation of the mark significantly more frequently during E5 (mirror, colour mark), than E2 (mirror, no mark), or E3 (mirror, transparent sham mark). Two independent and comprehensive analyses of the videos were conducted (by MK, JA), as well as two further blind analyses by unrelated researchers of a subset (15%) of the videos; the frequencies of the postures were highly correlated between the analyses (*r* = 0.988).

Posturing behaviours against the marked sites during periods E2 and E3 were infrequent and were not different between the two periods (Fig. 2Ca), a pattern driven by all fish except fish #7, which showed equal distributions of viewing angles (Table 3). This shows that the marking procedure itself had minimal effect on fish behaviour. In contrast, time spent posturing while viewing the marked sites was significantly higher in the coloured-(E5) versus no-(E2) and sham-marked (E3) periods (Fig. 2Ca), as predicted. This pattern held for all individuals except fish #2, regardless of the sites marked (Table 3). Note that no comparisons to E4 can be made with respect to observations of reflections, as no mirror was present during that period. Moreover, the time spent in postures reflecting the two remaining unmarked sites (e.g. right side of head and throat, for a fish marked on the left side of head) for each fish were not different among periods (Fig. 2Cb). Taken together, these findings demonstrate that cleaner wrasse spend significantly longer in postures that would allow them to observe colour-marked sites in the mirror reflection. These reactions also demonstrate that tactile stimuli alone are insufficient to elicit these behaviours, as they were only observed in the colour mark/mirror condition. Rather, direct visual cues, or a combination of visual and tactile stimuli, are essential for posturing responses in the mirror test. In previous studies on dolphins, similar patterns of activity were considered to constitute self-directed behaviour [5].

**Table 3.**
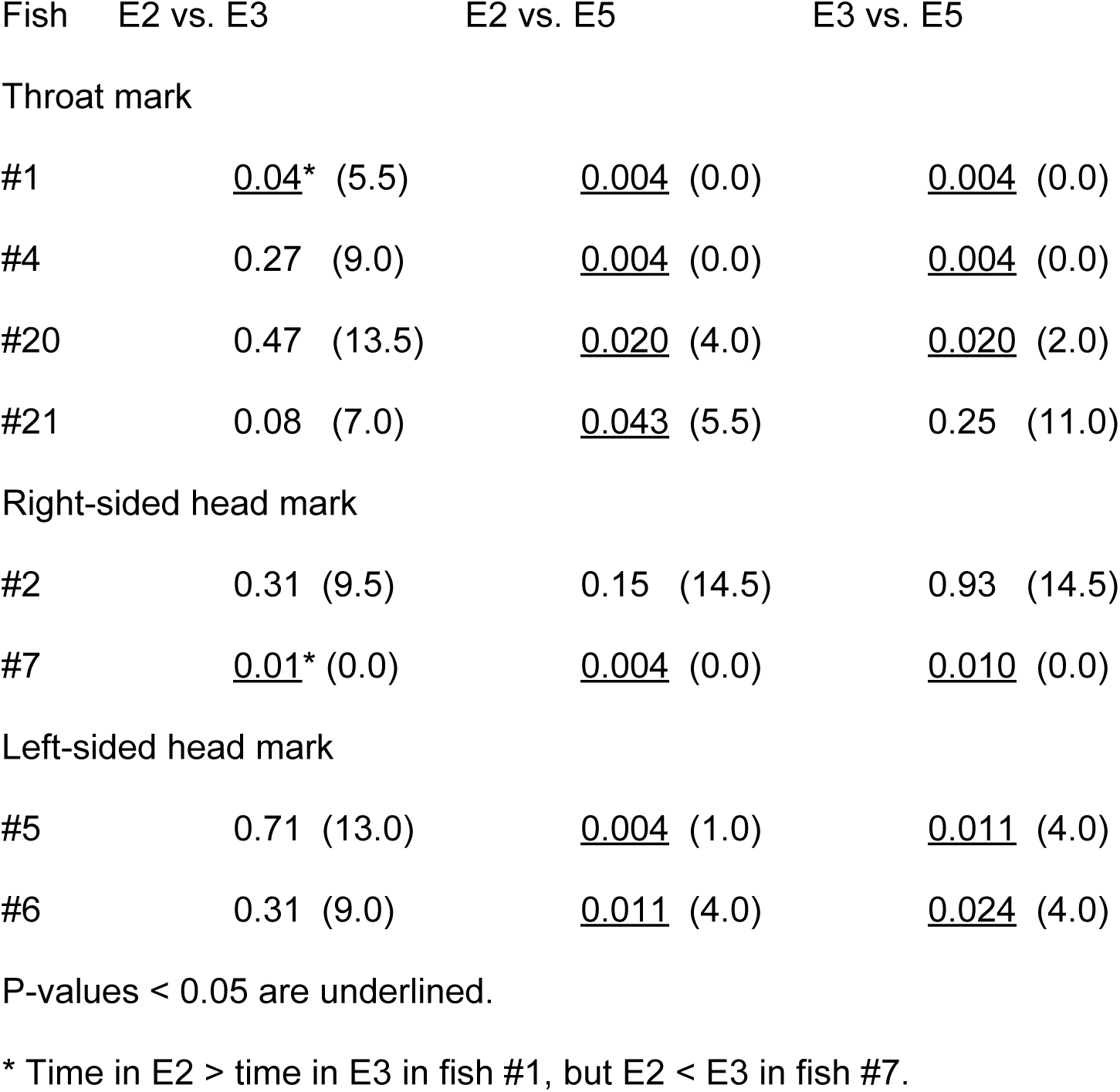
Comparison of time spent in postures in which the marked site was reflected between experimental periods of the mark test. Statistical tests (Mann–Whitney *U* test) at the level of the individual between E2 (no mark) and E3 (sham-mark), between E2 and E5 (coloured mark) and between E3 and E5. *P*-values, with the *U*-statistic in parentheses, are shown.

| Fish | E2 vs. E3 | E2 vs. E5 | E3 vs. E5 |
| --- | --- | --- | --- |
| 902 Throat mark |  |  |  |
| #1 | <u>0.04</u>* (5.5) | <u>0.004</u> (0.0) | <u>0.004</u> (0.0) |
| 904 #4 | 0.27 (9.0) | <u>0.004</u> (0.0) | <u>0.004</u> (0.0) |
| #20 | 0.47 (13.5) | <u>0.020</u> (4.0) | <u>0.020</u> (2.0) |
| 906 #21 | 0.08 (7.0) | <u>0.043</u> (5.5) | 0.25 (11.0) |
| Right-sided head mark |  |  |  |
| 908 #2 | 0.31 (9.5) | 0.15 (14.5) | 0.93 (14.5) |
| #7 | <u>0.01</u>* (0.0) | <u>0.004</u> (0.0) | <u>0.010</u> (0.0) |
| 910 Left-sided head mark |  |  |  |
| #5 | 0.71 (13.0) | <u>0.004</u> (1.0) | <u>0.011</u> (4.0) |
| 912 #6 | 0.31 (9.0) | <u>0.011</u> (4.0) | <u>0.024</u> (4.0) |

P-values < 0.05 are underlined.
914 \* Time in E2 > time in E3 in fish #1, but E2 < E3 in fish #7.

#### 2) Scraping of the colour-marked throat after viewing it in the mirror

Although they cannot touch their own bodies directly, many species of fish scrape their bodies on a substrate to remove irritants and/or ectoparasites from the skin surface [48,49]. When we marked fish with brown-pigmented elastomer on the lateral body surfaces in locations that could be viewed directly, the fish increased scraping behaviour of the mark sites, indicating they regard the colour dots as ectoparasites to be removed (Supplementary Figure S1). Similar scraping of colour-marked areas during the mark test is interpreted as an indicator of self-directed behaviour for some mammal species that do not have hands [29,50]. Accordingly, we hypothesised that the cleaner wrasse would scrape their bodies in an attempt to remove coloured marks from body parts not directly visible after observing them in the mirror (and crucially, that they did not scrape invisible sham marks, nor coloured marks in the absence of a mirror). As discussed earlier, if we observe a behaviour in a fish that is accepted to be functionally equivalent to a similar behaviour in mammals (in this case scraping), and that behaviour is accepted as being self-directed in those mammals [29, 50], then it raises the question whether this behaviour may be similarly considered self-directed in the fish. If this position is accepted, then any scraping behaviour of coloured marks in the mirror condition would constitute compelling evidence that fish use mark-directed behaviour to remove visually perceived coloured marks from their bodies. By extension and comparison to similar mirror test studies, this would raise the question of whether fish are therefore aware that the mirror reflection is a representation of their own body.

Like many natural behaviours, some scraping of the body flanks was observed outside the mirror condition in our studies. This body scraping behaviour was also difficult to distinguish from head scraping. Because of these factors we took throat scraping, and not face scraping, as the only evidence of a putative self-directed behaviour because it was never observed outside the period E5 in any of the subject fish. It is also important to note that fish marked on the head laterally scraped the body flank/facial region, but never the throat region, during period E5, providing further evidence that marking itself does not induce throat scraping.

Three of the four throat-marked fish frequently scraped their throats against the substrate after being exposed to the mirror during period E5 (Fig. 3A, B, Movies S4–S6), but none of the four fish exhibited this behaviour during E2–E4. We observed 37 separate instances of throat scraping during E5 (15 for fish #1, 16 for fish #4, 6 for fish #21; Friedman test, *χ*^2^ = 9.0, *df* = 3, *P* = 0.029; binomial test within individuals, E2, E3 and E4 vs. E5: 0 vs. 15 scrapings, *P* < 0.0001 in fish #1, 0 vs. 16 scrapings, *P* < 0.0001 in fish #4, 0 vs. 6 scrapings, *P* = 0.031 in fish #21). These three fish attempted to scrape their throats but this was occasionally executed awkwardly, probably because they were not accustomed to performing this behaviour. As the marks were identical in periods E4 and E5, with the only change being the visibility of the mirror, the difference in throat scraping provides further strong evidence that the colour injection itself did not cause direct physical stimulation that would lead to the observed behaviours (e.g. itching or pain).

**Fig. 3.**
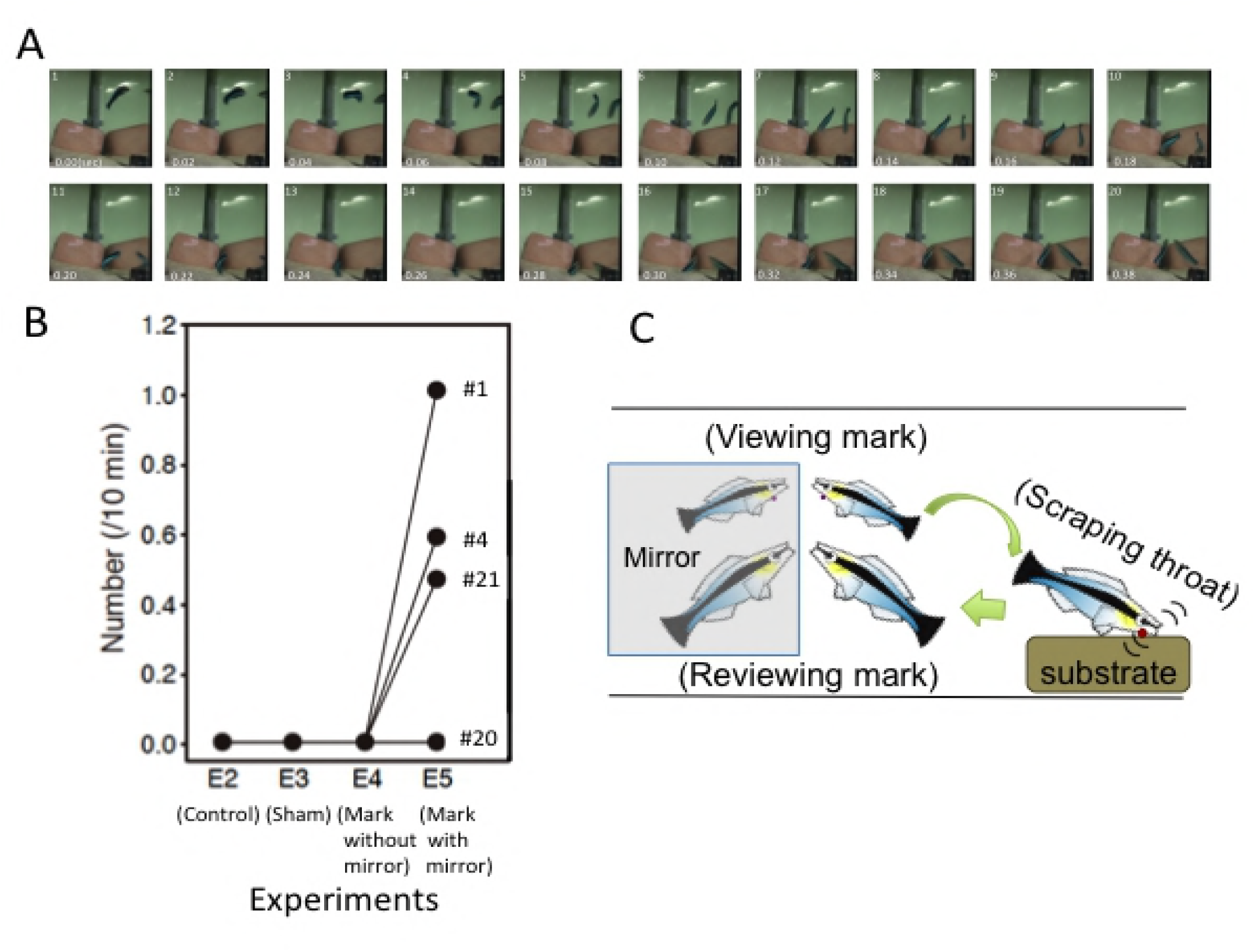
Throat scraping behaviour (self-directed behaviour) and its pattern of occurrence. (A) Frame-by-frame example of throat scraping behaviour. (B) Frequency of throat scraping behaviour of the four throat-marked fish during periods E2, E3, E4 and E5. (C) Schematic sequence of throat scraping behaviour including viewing, and then reviewing, the throat (before and after scraping, respectively).

These results accord with the increased amount of time spent in postures indicating observation of the coloured marks in the reflection only during period E5 (Fig. 2Ca). The motivation for scraping the mark is potentially to remove a perceived ectoparasite, which these wild-caught fish would have experienced previously. In all cases (*n* = 37), the scraping behaviours followed soon after the fish had assumed a posture that reflected the throat mark, with an average latency between observation in the mirror and scraping of the substrate of 1.93 sec ± 1.16 (*n* = 37; see Fig. 3C, Movies S4–S6). However, posturing was not always followed by scraping. The physical substrate on which fish scraped varied among individuals: all scraping was done in a narrow area of the sandy bottom by fish #1, and all and the majority (14/16) of the scraping of fishes #21 and #4 was done on a rock in the corner of the tank despite the same arrangement in all tanks. This may be because fish learn successful scraping techniques associated with specific substrates and continue to use them.

The majority of the throat scraping behaviour was immediately followed by another frontal-vertical posture performed in front of the mirror (after 1.82 sec ± 1.46; *n* = 31; Movie S6). Assuming frontal-vertical postures during swimming is atypical, and in these postures observation of the marked site via the mirror-reflection is possible. As such, this pattern of behaviour may constitute contingency testing, in this case to check if the perceived parasite had been removed by the scraping attempt. While this interpretation does imply intentionality on the part of the animal, the general rarity of this behavioural sequence, and frequency with which it was displayed during the mark test, provide compelling evidence for this interpretation. Indeed, this type of behaviour is similar to that of chimpanzees, which examine and smell their fingers after touching a paint mark [1,8], and which is considered intentional self-directed behaviour in that species.

Three of the four throat-marked individuals in this study passed the mark test, a success ratio comparable to other species tested previously; one of three Asian elephants passed the test [4], as did two of five magpies [6]. Fish #20 in our study was throat-marked but did not perform throat scraping (Fig. 3B). However, this fish exhibited intensive contingency-testing behaviours (*a*) and (*b*) during period E1, prior to colour-marking, similar to the other fish (Table 2), and assumed postures (self-directed behaviour) that reflected the throat more frequently during E5 after colour marking (Table 3). According to the mark test criteria used for dolphins [5], these results suggest that this wrasse recognised the reflection as self, but “fell at the last hurdle”. Nevertheless, given the controversial nature of the mark test in non-primates, and questions over the interpretation of these behaviours [8], we do not take this result as conclusive evidence for MSR in this individual. We do point out, however, that by the same criteria used for e.g. dolphins, we would conclude that all four throat-marked fish recognised themselves in the mirror.

In this study we applied the mark test, a controversial assessment of animal cognition [8], to a fish, a taxonomic group often considered to have lower cognitive abilities than other vertebrate taxa. We find compelling evidence that cleaner wrasse pass through all stages of the mark test, ultimately attempting to remove the mark when it is able to be viewed in the mirror (Figure 3). We further find the parsimonious conclusion to be that the behaviours displayed by this fish are equivalent to behaviours taken as evidence for self-recognition in other taxa (contingency testing, self-directed behaviour, observation and exploration of the body in a reflected image, and removal attempts; Figures 1,2,3). We consider these behavioural responses to be a consequence of the particular feeding ecology, generally high cognitive capacity, and problem-solving skills of the cleaner wrasse [14-16,37,38]. This is the first report of successful passing of the mark test in vertebrates outside of mammals and birds, suggesting that if mirror tests are applied considering the cognitive capacities and ecology of focal species outside of primates, they too may pass the test. Our study further supports previous theories postulating that recognition and cognitive capacities are more closely related to social and behavioural ecology than relative brain size or phylogenetic proximity to humans [14,16,32].

The results we present here will by their nature lead to controversy and dispute, and we welcome this discussion. We consider three possible interpretations of our results and their significance for understanding the mark test: i) the behaviours we document are not self-directed and so the cleaner wrasse does not pass the mark test, ii) cleaner wrasse pass the mark test and are therefore self-aware, or iii) cleaner wrasse do pass the mark test but this does not mean they are self-aware. If one takes position i), rejecting the interpretation that these behaviours are self-directed, it is necessary to demonstrate grounds for this rejection. As noted above, touching or scraping behaviour is taken as evidence of a self-directed behaviour in mammals, and so if these behaviours are not similarly considered self-directed in fish, the question must be asked why. For a test to be applicable across species, an objective standard is required. Without such a standard, behaviours assessed in the mark test can be differently assessed depending on the taxon being investigated. This introduces an impossible, and unscientific, standard for comparison and we therefore reject this conclusion or must reject the validity of the mark-test entirely.

We therefore consider the most parsimonious conclusion to be that the behaviours we observe here in cleaner wrasse are equivalent to those in other taxa during the mirror test. Based on this, and on the original interpretation of the mark test by its inventor Gallup, who suggested species that pass the mark test are self-conscious and have a true theory of mind [1,57], would therefore lead us to take position (ii), that cleaner wrasse are self-aware. However, we are more reserved about the interpretation of these behaviours during the mark test with respect to self-awareness in animals. We do not consider that the successful behavioural responses to all phases of the mark test should be taken as evidence of self-awareness in the cleaner wrasse, but rather that these fish come to understand that the mirror reflection represents their own body. From the behaviour we observe, we consider the interpretation that makes fewest assumptions to be that these fish undergo a process of self-referencing, whereby the fish use the mirror to see their own body, but without this involving theory of mind or self-awareness [32]. This interpretation is supported by a supplementary experiment (Supplementary Figure S1) that showed fish marked on the body in places they could directly see also performed scraping on those regions.

If we therefore accept position (iii), that cleaner wrasse show behavioural responses that fulfil the criteria of the mark test, but that this result does not mean they are self-aware, a question naturally arises. Can passing the mark-test be taken as evidence of self-awareness in one taxon but not another? A position that holds the same results can be interpreted different ways depending on where they are gathered is logically untenable, and so must be rejected. This leaves us with the only option to re-evaluate our interpretation of what the mark-test means, in particular to reject the position that successfully passing the mark test means animals are self-aware and accept that successful performance in the mark test may be driven by numerous processes. To hold any other dualistic interpretation of the test would be taxonomically chauvinistic and would undermine the standing of the mark test as a valid metric of self-cognizance in animals (34). Based on our findings, we therefore advocate for a reappraisal of the interpretation of the mark test, and conclude that many more species may be able to pass the test when it is applied in a manner that is sympathetic to their natural biology.

## MATERIAL AND METHODS

### Animals and housing

The cleaner wrasse, *Labroides dimidiatus*, is a protogynous hermaphrodite that lives in coral reef habitats [46,58]. We used 10 wild fish obtained from commercial collectors in this study. Prior to our experiments, the fish were housed in separate tanks (45 × 30 × 28 cm^3^, Fig. 1A) and each fish was kept for at least 1 month prior to beginning the experiments to ensure acclimation to captivity and the testing conditions, and that they were eating and behaving normally. Fish were between 51–68 mm in length; this is smaller than the minimum male size, thus strongly suggesting that these individuals were functionally female. Individual fish sizes were as follows: 68 mm for fish #1, 62 mm for fishes #13 and #20, 61 mm for fish #21, 58 mm for fish #4, 55 mm for fish #5, 53 mm for fish #6, 52 mm for fish #2, and #7 and 51 mm for fish #3). Each tank contained a 5 × 5 × 10 cm^3^ rock in the corner, and a PVC pipe that provided shelter on a coral-sand substrate 3–4 cm deep. The water was maintained at 24–26°C and was aerated and filtered. The fish were maintained on a 12 h: 12 h light: dark cycle. Artificial flake food (Tetramin; Tetra, Blacksburg, VA, USA) and small pieces of diced fresh shrimp were given twice daily.

### Mirror presentation to fish

The mirror presentation method (e.g. duration, timing, position and mirror size and shape) has important consequences for successful implementation of MSR studies [1,4,5]. We presented a 45 × 30 cm^2^ high-quality mirror on a glass wall of the same size inside the experimental tank (Fig. 1A). The mirror was positioned at night, while the fish were sheltered within the PVC pipe, 1 week before beginning the experiments; it was then completely covered with a white plastic sheet (45 × 30 cm^2^). At the start of the experiments, the white cover on the mirror was removed, and the subject fish were exposed to the mirror until the end of the series of experiments, with the exception of a half-day experiment during which the mirror was completely covered with the white sheet (see below).

### Order of presentation of the five experiments, E1–E5

We studied fish behavioural responses during five experimental periods, from Experiment 1 (E1) to Experiment 5 (E5) (in chronological order). All behaviours were recorded by video camera (HDR-CX370; Sony, Tokyo, Japan), and the field of view encompassed the entire experimental tank (Fig. 1A). After the start of the experiment (i.e. uncovering of the mirror), the behavioural responses of fish were video-recorded eight times over a 2-week period: every day for the first 5 days, and then again on days 7, 10 and 15 (period E1: initial responses to mirror). Three or four days after the end of period E1, fish behaviours were recorded for 2 consecutive days as a control for the mark test (E2: no mark). Two or three days after E2, a non-visible sham-mark (white or non-pigmented elastomer) was made on all 10 fish, and behavioural responses were recorded over the next 2 days (period E3: sham-marking). Two days after period E3, the fish were marked with a coloured (brown) elastomer mark at night, and their behaviours were recorded in the morning of the following day while the mirror was covered with the 45 × 30 cm^2^ white plastic sheet (period E4: colour mark with no mirror present). After this observation phase, the mirror was uncovered and behaviours were recorded on the afternoon of the same day, and also on the morning of the following day (period E5; colour-mark with mirror present). All experiments were necessarily run in this order because coloured tags could not be removed from fish once applied; hence, transparent (sham) marks always preceded coloured marks. The four fish that were marked on the head showed an increase in scraping of the marked area during period E5. However, three of these fish were also observed scraping facial areas prior to colour marking, indicating that face-scraping cannot be taken as unequivocal evidence of mirror-induced self-directed behaviour.

### Provisioning mark procedure

Elastomer tags and visible implant elastomer (VIE) marking (Northwest Marine Technology Inc., Shaw Island, USA) via subcutaneous injection are widely used in studies of individually marked live fish and do not affect fish behaviour [51-54, NMTI]. Our fish were taken from their tanks at night together with their PVC pipe, and placed in eugenol solution to achieve mild anaesthesia (using FA100; Tanabe Pharmacy Inc., Japan). An un-pigmented gel mark was injected subcutaneously in an area of 1 × 2 mm^2^ at one of three sites during the sham mark period: on the right side of the head (two fish), on the left side of the head (two fish) or under the throat (four fish; Fig. 2A). The entire injection process took no longer than 5 minutes, and the fish were returned to their original tank together with the pipe after the mirror was covered with the white plastic sheet. We ensured that the fish were swimming normally the next early morning, and showed no behavioural changes as a consequence of the tagging procedure. We initially used white pigment on the pale-coloured body areas, but found that the skin in these areas had a slight blue tint, and that the white tag was visible in two fish; these fish were not used in further experiments. A brown-pigmented elastomer colour mark was applied as colour-mark at night before the day of E4. After confirming that all marks were of the same size (1 × 2 mm^2^), the fish were returned to the tank. Given the location of the tags relative to the field of view of cleaner wrasse, direct observation of the marks on the head was unlikely, and was definitely impossible for throat marks. To standardise the testing procedure, the brown-coloured mark was injected at the throat near the transparent marked site. Even with both marks applied, the total volume of the tag was lower than the minimum recommended amount, even for small fish, and < 13% of the size of tags used in studies with other fish: biologists who applied VIE to small fish in previous studies, i.e. 26-mm brown trout [51] and 8-mm damselfish [54] stated that the amounts used were minute, but for the former species 2–3 mm tags were made with 29 G needles [51]. Willis and Babcock used large tags (10 ×1 × 1 mm (127/ml) in *Pagrus auratus* (from NMTI) [53]. Our own tagging method was therefore very unlikely to have caused irritation. Moreover, we saw no evidence during period E4 (colour tag, no mirror present) of any removal attempts or scratching behaviour, further confirming that the tags did not stimulate the fish.

### Behavioural analyses

Videos of the fish behaviours were used for all behavioural analyses. Fish performed mouth-to-mouth fighting frequently during period E1, and the duration of this behaviour was recorded (Fig. 1B, Movie S1). Unusual behaviours performed in front of the mirror, which have never been observed before in a mirror presentation task, nor in the presence of a conspecific, were often observed during the first week of E1, and the type and frequency of these behaviours was recorded.

### Description of postural behaviours performed in front of the mirror and behavioural observations

In the latter half of E1, fish occasionally swam slowly or remained stationary in front of the mirror, and the duration (in seconds) of these behaviours, when performed within 5 cm of the mirror, was recorded. The duration of postures in which the marked area was reflected in the mirror (i.e. viewing behaviours) was recorded during E2 (no mark), E3 (sham mark) and E5 (coloured mark with mirror present). Posturing within 5 cm of the mirror was categorised into three types: right sided posture (i.e. reflecting the right side of the head), left-sided posture (reflecting the left side of the head) and frontal-vertical posture (reflecting the throat). The duration (in seconds) of each of the three types of posture was recorded during six separate 5-min observation periods, for a total of 30 min per fish for each of the periods when a mirror was present (E2, E3 and E5). A subset of 15% of the videos was blindly analysed by two researchers outside our team; their analysis was highly correlated with the main analysis (r = 0.887, *P* < 0.0001), and statistical tests showed no significant differences between the two datasets (two-way repeated-measures analysis of variance [ANOVA], blind effect: *F* = 0.06, *P* = 0.80, blind effect × observation site: *F* = 0.77, *P*= 0.45).

Scraping behaviour, including the location on the body that was scraped, was observed during periods E2–E5 in the eight subject fish. During period E5, when the fish were colour-marked and exposed to the mirror, individuals often displayed the marked site to the mirror immediately prior to and following a scraping behaviour. Therefore, we also recorded the time interval between displaying and scraping during E5.

### Responses towards real fish

A potential alternative explanation of behaviour in mark tests (and one that is rarely tested for in other vertebrates) is that the focal individual perceives their reflection not as the self, but rather as another individual behind a glass divide. Although many behaviours seen in the mark test suggest that this is not the case (e.g. contingency-testing, body exploration), and a growing body of evidence shows that fish perceive mirror reflections in a fundamentally different way to conspecifics behind glass [59,60], we directly controlled for this possibility by comparing the behaviour of fish confronted with a reflection to that when another individual was across a glass divide.

We tested the responses of eight fish (55–59 mm in size) in size-matched pairs. Two fish were introduced into a tank (45 × 30 × 26 cm^3^). After the fish became acclimated, the cover was removed from the divider to allow them to see one another. We then recorded behavioural responses in the same manner as described for period E1 in the mirror test, for 2 weeks. The results of these observation are presented in Fig. 1D. After 3 weeks, we marked these fish on the throat and recorded whether they scraped their throat regions; however, we did not observe any throat-scraping behaviour, although they must have observed the ‘parasite’ on the throat of the conspecifics. This indirectly supports the view that the fish were attempting to remove the mark from their own bodies when presented with the mirror during period E5 of the actual mark test.

### Statistical analyses

Statistical analyses were performed using SPSS (ver. 12.0; SPSS Inc., Chicago, IL, USA) and R software (ver. 2.13.2; R Development Core Team 2011). During period E1, the responses of the subject fish to the exposed mirror changed significantly over time. Changes in the duration of mouth fighting and time spent within 5 cm of the mirror over time were analysed with linear mixed models (LMMs). Similarly, changes over time in the duration of mouth fighting and time spent within 5 cm of the mirror were analysed with LMMs for the experiments using real fish across glass dividers. The frequency of unusual mirror-testing behaviours was analysed using a generalised linear mixed model (GLMM) with a log-link function and assuming a Poisson distribution. Time spent in postures reflecting the right side of the head, left side of the head and the throat were compared between mark types during the mark tests (E2: no mark, E3: sham mark and E5: coloured mark with mirror present) using repeated-measures ANOVA. Note that the marked and unmarked positions were analysed separately (Fig. 2Ca, b). Individual-level statistics on postures that reflected the marked sites are shown in Table 3 (Mann–Whitney *U* test with duration in seconds of the six different behaviours per 5-min observation in periods E2, E3 and E5). To detect the effect of throat marking on the frequency of scraping behaviour, a Friedman test was used on the entire data set (E5 vs. E2, E3 and E4) and a binomial test was used for comparison between periods (E5 vs. E2, E3 and E4). No throat scraping or unusual behaviours were observed when individuals interacted with conspecifics across a glass divider, so no statistical tests were performed for that condition.

### Ethics statements

Our experiments were conducted in compliance with the animal welfare guidelines of the Japan Ethological Society, and were specifically approved by the Animal Care and Use Committee of Osaka City University.

## SUPPORTING INFORMATION

### Responses to color dot on flank directly visible and color dot on the mirror

If cleaner fish pass the mark-test, they will also scrape the mark when they detect directly it on its flank without the aid of mirror. We examined whether cleaner wrasse respond to the mark dot on the flank without mirror. Five other fish of the same size range were kept in experiment tanks without mirror, and their behaviours were video-recorded for 3 hours in each three conditions in the chronological order of, i) no marking as control, ii) transparent sham marking on the center of left body side, directly visible for the fish, and iii) colored marking on the left area of the same side of flank, using the same procedure of the mark test. The results are shown in Supplementary Figure S1. Both sides of the body were scraped on substrate infrequently in the control i) and sham mark ii). In contrast, in the marked phase iii), the marked site in left flank were frequently rubbed more than right side in the same phase, and than left side of control and sham marking phases (Interaction: *χ*^2^ = 12.35, *df* = 2, *n* = 5, *P* < 0.002), strongly indicating that cleaner fish regard the directly visible color dot on body surface as ectoparasite.

Cleaner wrasse will regard the color dots on body skin as ectoparasite, but do they also regard the same dots on elsewhere, e.g. on mirror as the ectoparasite? We video-recorded and examined their responses of the other 4 fish to the color dot on mirror on the first day and 10 days after mirror presentation. On 10 days fish will do MSR. The video-record of 30 min showed all fish ignored, hardly watched and approached the color dots on the mirror, and never stayed in front of it in both days, indicating fish did not pay attention to the dot. Cleaner fish show directed removal attempts toward the mark (Supplementary Figure S1), indicating that marks on the body are perceived differently to those elsewhere in the environment, whether these bodily marks are observed directly or with a mirror.

### Visual and tactile stimuli by the colour mark

We further considered whether the elastomer tag could provide a tactile stimulus that, when paired with visual information, may lead to individuals passing the mark test [8,55]. We can effectively rule out that tactile stimulation alone was sufficient to induce a self-directed behaviour [55] because we observed no throat scraping in any fish during periods E3 (sham mark) and E4 (coloured mark with no mirror present). This is in contrast to previous studies on rhesus monkeys using a somatosensory training stimulation in which the stimulus immediately elicited a response due to irritation [10,11] – we observed no such spontaneous response to the tactile stimulus. Moreover, and likely due to the inherent tendency of the species to search for ectoparasites on the body surface of clients and attempt to remove them [37,45], we did not observe a temporal progression in behaviour that may suggest direct or indirect association learning [8,10,56], rather a rapid onset of removal attempts in period E5. We therefore conclude that the behaviours we observe were primarily driven by visual stimulation, and required no association learning, but acknowledge the provision of elastomer tags may provide more tactile stimulation than paint marks [e.g. 1,4]. Future studies should attempt to experimentally disentangle these two stimuli to assess the roles of visual and tactile stimulation in mirror test with the cleaner wrasse.

## Acknowledgements

We are grateful to numerous conference attendees and anonymous reviewers for their constructive and vigorous advice on the experiments and the manuscript, and to members of the Laboratory of Animal Sociology, Osaka City University for their support and fruitful discussions. Financial support by a Grant-in-Aid for Scientific Research from the Ministry of Education, Culture, Sports, Science and Technology (MEXT), Japan (to MK), and by Japan Society for the Promotion of Science (to LAJ) is gratefully acknowledged.

## Author contributions

MK, TT and LAJ conceived and designed the experiments. MK, TT, TH, HT and JA performed the experiments. SA and HT analysed the data, MK and LAJ wrote the manuscript.

## SUPPLEMENTARY MATERIALS

One figure and six movies of captured cleaner wrasse behaviours during the mirror tests, i.e. aggressive fighting, idiosyncratic mirror-testing behaviour, and scraping of coloured marks on the throat.

**Supplementary Figure S1.** The frequencies of rubbing body sides by cleaner wrasse before marking (Control), after transparent marking (Sham) and colour marking (Mark) during 3 hours without mirror. Sham and colour marking were on left flank, the area directly visible for fish (*χ*^2^ = 12.35, *df* = 2, *n* = 5, *P*<0.002)

## Supplementary Movies

**Movie S1**. Mouth-to-mouth fighting against the mirror reflection on the second day after the initial mirror presentation in phase (*i*) (fish #1). The fish attack the reflection with open mouths during fighting in a common display of fish aggression.

**Movie S2**. An example of idiosyncratic behaviour. Mirror-testing behaviour (upside-down approach) performed by fish #1 on day 4. The fish approached the reflection in an upside-down position, but returned in front of the mirror and viewed the mirror following the behaviour.

**Movie S3**. An example of idiosyncratic behaviour. Mirror-testing behaviour (dashing along mirror; ‘rapid dash’) performed by fish #21 on day 4. The fish did not attack the mirror reflection, but looked at the mirror following the behaviour.

**Movie S4**. Scraping of the throat mark on the sandy bottom by fish #1 immediately after viewing the mark in the mirror. The fish assumed a position that reflects the throat in the mirror soon after scraping.

**Movie S5**. Fish #1 tried to scrape a throat mark on the sandy bottom immediately after viewing the mark in the mirror, but did not look at its throat in the mirror after scraping. However, the fish failed to scrape its throat on the sandy bottom, although the sand moved as the fish shook its head. The fish may not have checked its throat in the mirror, possibly because it had not been scraped.

**Movie S6**. The fish rapidly approached the mirror after scraping its throat on the sandy substrate, stopped at a distance of about 1 cm from the mirror, and remained stationary for 1 s; during this time the fish assumed a position that reflected the scraped throat in the mirror.

---

Table 1: Description of five types of atypical (mirror- or contingency-testing) behaviours frequently observed during days 3–5 after presentation of the mirror.

*(a)* Dashing along the mirror: rapid dashing along the mirror surface in a single direction for 10–30 cm. Fish did not swim directly against or make contact with their mirror reflection.

*(b)* Dashing along the mirror with the head in contact with the mirror: the head of the fish was always in contact with the mirror during dashing.

*(c)* Dashing and stopping: fish rapidly dashed towards the mirror reflection but stopped before contact with the mirror.

*(d)* Upside-down approach: fish swam in an upside-down posture while approaching the mirror. *(e)* Quick dance: fish spread all of their fins, and quickly arched and quivered the body several times during ca. 1 s at a distance 5–10 cm from the mirror; no dashing to the mirror was observed.

